# Acoustic biomolecules enhance hemodynamic functional ultrasound imaging of neural activity

**DOI:** 10.1101/734939

**Authors:** David Maresca, Thomas Payen, Audrey Lee-Gosselin, Bill Ling, Dina Malounda, Charlie Demené, Mickaël Tanter, Mikhail G. Shapiro

**Affiliations:** Division of Chemistry and Chemical Engineering, California Institute of Technology, Pasadena, CA, USA; Physics for Medicine Paris, INSERM, CNRS, ESPCI, Paris, France

## Abstract

Hemodynamic functional ultrasound imaging (fUS) of neural activity provides a unique combination of spatial coverage, spatiotemporal resolution and compatibility with freely moving animals. However, deep and transcranial monitoring of brain activity and the imaging of dynamics in slow-flowing blood vessels remains challenging. To enhance fUS capabilities, we introduce biomolecular hemodynamic enhancers based on gas vesicles (GVs), genetically encodable ultrasound contrast agents derived from buoyant photosynthetic microorganisms. We show that intravenously infused GVs enhance ultrafast Doppler ultrasound contrast and visually-evoked hemodynamic contrast in transcranial fUS of the mouse brain. This hemodynamic contrast enhancement is smoother than that provided by conventional microbubbles, allowing GVs to more reliably amplify neuroimaging signals.

## INTRODUCTION

Functional ultrasound imaging (fUS) is a breakthrough technology that uses ultrafast frame rates to map changes in local cerebral blood volume induced by neural activity (1). Due to its high spatiotemporal resolution and versatile form factor, fUS has emerged as an attractive basic neuroscience tool capable of visualizing whole-brain functional activity in a variety of animal models and humans (2), providing functional resolution of the order of 100 μm (3, 4).

Unfortunately, skull bones attenuate and distort ultrasound waves in the MHz range (5), which hinders the fully noninvasive potential of fUS imaging. As a result, except for a few studies (6), the vast majority of fUS imaging has been conducted using craniotomies (1, 4, 7, 8), thinned-skull preparations (9), or acoustically transparent windows (2, 10).

A potential solution to compensate for skull attenuation is to augment the source of contrast in fUS – red blood cells – by administering ultrasound contrast agents into the blood stream. Errico et al. (9) showed that commercial lipid-shelled microbubbles (MBs) can enhance transcranial hemodynamic signals in rats. However, despite clear improvements in signal, this approach suffered from several limitations related to the physics of MBs. First, as MB-backscattered intensity decays rapidly over time due to gas diffusion (11), organ retention (12) and MB deflation upon ultrasound exposure (13), it was necessary to administer 13 bolus injections of MBs in rats over the course of 10 minutes (9), resulting in an exceptionally high dose. A second limitation is that MBs add significant random fluctuations to fUS signals (see (9), and Fig. 3F), which lowers the correlation score of functional activity maps. These fluctuations are likely due to the variability in the acoustic response of MBs of a given size (14), the lipid shell surface tension (15), the agent polydispersity (diameters ranging from ~ 1 μm to >10 μm), and the pressure-dependent attenuation and scattering of MBs (16).

Here, we introduce a new class of hemodynamic enhancers for fUS based on acoustic biomolecules known as gas vesicles (GVs) (17, 18). GVs comprise air-filled compartments with dimensions on the order of 200 nm, enclosed by a 2 nm-thick protein shell. These nanostructures were recently introduced as biomolecular ultrasound contrast agents and acoustic reporter genes (19–22). We hypothesized that the physical properties of GVs would provide potential advantages for hemodynamic enhancement. GVs are physically stable, relatively monodisperse in their cylindrical diameter, and much smaller than MBs. Thus, at comparable gas volume fractions these more numerous, stable, and monodisperse contrast agents are expected to boost hemodynamic contrast with substantially less fluctuation. Here we tested this hypothesis by examining the fUS enhancement provided by GVs purified from *Anabaena flos-aquae* (23), which in their wild-type form serve as non-resonant linear scatters at biomedical ultrasound frequencies (1-30 MHz) (24). We compared these biomolecular contrast agents to commercial MBs *in vitro* and *in vivo*.

## MATERIAL AND METHODS

### Gas vesicle preparation

Gas vesicles were isolated from *Anabaena flos-aquae* via hypertonic lysis and buoyancy purification using previously described protocols (23). Their concentration was measured using their optical density at 500 nm (OD_500nm_).

### Flow phantom design

To estimate the flow detection limit of GV-enhanced fUS imaging, we prepared three model solutions containing subwavelength ultrasound scatterers (Fig. 1A). The first solution was a commercial Doppler fluid (DF) (Model 707, ATS Laboratories, Bridgeport CT, USA) containing linear particles mimicking red blood cell scattering. The second solution contained the DF mixed with a commercial MB contrast agent (SonoVue, Bracco Imaging, Geneva, Switzerland) diluted at a ratio of 1:1000 (5 × 10^5^ microbubbles per mL). The third solution contained the DF mixed with purified gas vesicles (23) at a concentration of 10^11^ GVs per mL. Each solution was flowed through a 1.5 mm diameter channel extruded in a tissue-mimicking ultrasound phantom (1% cellulose embedded in 5% agar solution) using a syringe pump (GenieTouch, Kent Scientific Corportion). The channel and the surface of the phantom both formed a 5° angle with respect to the ultrasound probe (Fig. 1B) in order to prevent attenuation heterogeneity due to different phantom thickness above the channel.

**Figure 1.**
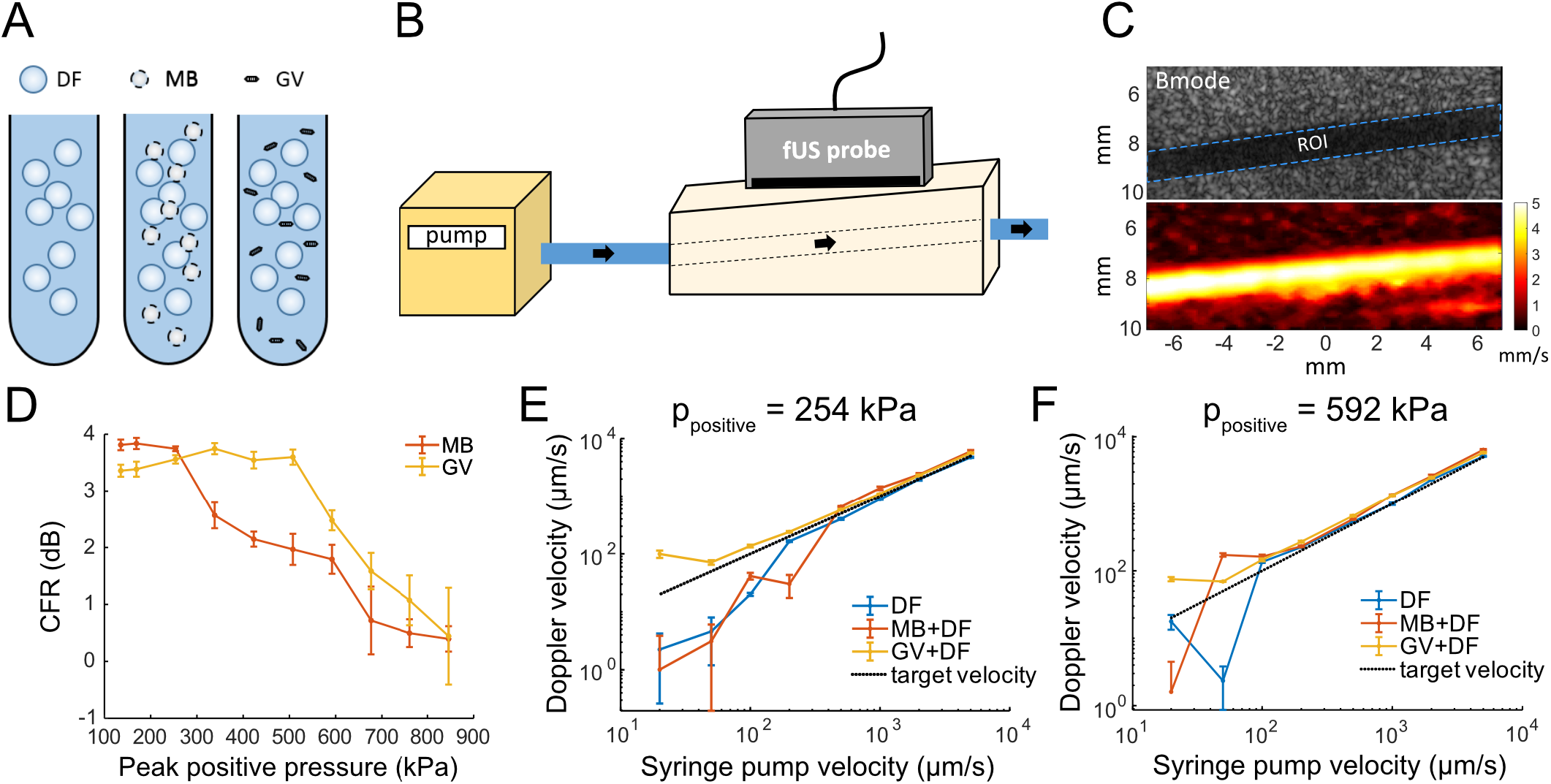
GVs estimate slow flows more accurately and sustain higher pressure than MBs. A/ The three contrast enhancing solutions investigated: Doppler fluid (DF), microbubbles (MBs) and gas vesicles (GVs). B/ Schematic of the flow phantom setup. C/ Conventional ultrasound B-mode image indicating the selected ROI, and corresponding power Doppler image of the flow phantom. D/ Contrast-to-Doppler-fluid ratio (CFR) as a function of pressure at a flow velocity of 2 mm/s. E/ Flow measurement correspondence of DF, GV, and MB-enhanced ultrafast Doppler imaging at 254 kPa. F/ Flow measurement correspondence of DF, GV, and MB-enhanced ultrafast Doppler imaging at 592 kPa.

### Functional ultrasound imaging sequence

Ultrasound imaging was performed at 15 MHz using a 128-element linear probe (Vermon, Tours, France) connected to a programmable ultrasound scanner (Verasonics Vantage, Seattle, USA). We transmitted angled ultrasound plane-waves (−6:2:6 degrees) at a 7 kHz framerate, which resulted in a 1 kHz framerate after coherent compounding (25). Ensembles of 200 compounded frames were acquired every 0.5 s and processed using a singular value decomposition (SVD) filter (26) to generate power Doppler images (27) of the flow phantom (Fig. 1C) or of the mouse brain (Fig. 2B), resulting in a 2 Hz fUS imaging framerate.

**Figure 2.**
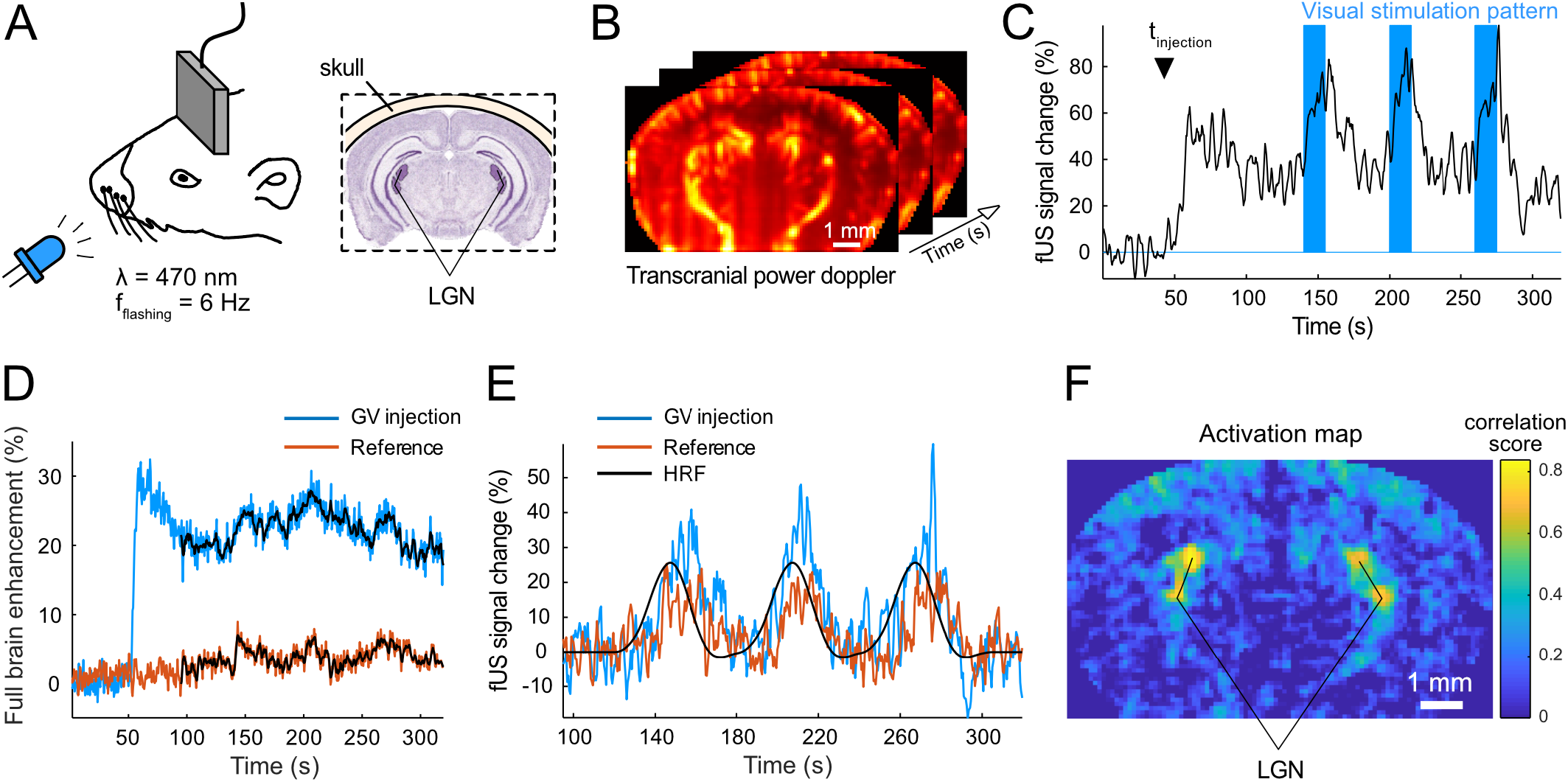
Visual stimulation protocol. A/ Illustration of a lightly anesthetized, head fixed wild type mouse exposed to a flashing blue light. Probe positioned to capture the lateral geniculate nuclei (LGN). B/ Transcranial ultrafast Doppler image acquired at 2 Hz over the course of 320 seconds. C/ Visual stimulation pattern (blue) and corresponding fUS signal response in the LGN in the presence of GVs (black). D/ Mean brain enhancement with and without GVs. E/ Truncated fUS signals in the most responsive pixel with and without GV injection. The predicted hemodynamic response function (HRF) is overlaid on the experimental activation patterns. F/ fUS activation maps revealing activation of the LGN.

In the flow phantom study, we selected an ultrasound imaging plane displaying the longitudinal cross section of the phantom channel (Fig. 1C), which was positioned at the geometric focus of the ultrasound probe (8 mm). For *in vivo* experiments, we selected manually the plane of interest that contained the lateral geniculate nuclei (LGN) – a sub-cortical structure of the mouse visual system (Fig. 2A) – and positioned the outer skull surface at a 2.5 mm distance from the ultrasound probe surface.

### Flow phantom data processing

To quantify fUS signal enhancement for each of the contrast agent solutions injected in the phantom channel, we manually selected a region-of-interest inside the channel (ROI) as illustrated in Fig. 1C. Each data point shows the mean±SEM resulting from 30 measurements. First, we investigated the resistance of the different solutions to ultrasound pressure in these flow phantom conditions. Each solution was injected at 2 mm/s and insonated with peak positive pressures ranging from 135 kPa to 2.1 MPa. We calculated the contrast-to-Doppler-fluid ratio (CFR) in decibels (dB) by measuring the mean power Doppler intensities 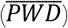 in the ROI for the ultrasound contrast agent (UCA) and the Doppler fluid (DF) respectively, according to the following formula:

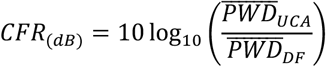

Then, using the syringe pump, each scattering solution was circulated in the phantom channel at flow velocities ranging from 5 mm/s to 20 μm/s and insonated at 254 kPa and 592 kPa. The corresponding Doppler-derived flow velocities were extracted by retrieving the Doppler frequency f_Doppler_ from the phase of the signal, and from the knowledge of the channel angle θ relative to the ultrasound propagation direction, according to the following classic formula:

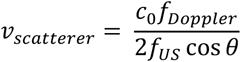

where *c*_*0*_ is the speed of sound in the medium of interest, and *fUS* is the transmitted ultrasound frequency.

### Animal preparation

All *in vivo* experiments were approved by the Institutional Animal Care and Use Committee of the California Institute of Technology. C57BL/6J male mice aged between 9 and 11 weeks were used in this study. Mice were anaesthetized using 1–2.5% isoflurane using a nose cone, depilated over the head and placed on a 37 °C heating pad. A catheter was inserted in the tail vein for injection and affixed in place using GLUture. Mice were then anesthetized with Ketamine/Xylazine and placed on a stereotaxic frame for the imaging session. Ultrasound gel, previously centrifuged at 2000 G for 10 minutes to remove bubbles, was applied to couple the transducer probe to the animal. 45 s after the beginning of the imaging sequence, 50 μL of saline, GVs (OD_500nm_=100, corresponding to 11.4 nM) or Definity MBs (Lantheus Medical Imaging, N. Billerica, MA, USA) diluted to 10^8^ microbubbles/mL were injected intravenously using the catheter. The rate of injection was 5 μL/s.

The catheter was made from PE10 tubing and a 30 G needle. MBs were handled according to the manufacturer’s instructions and passed through a 30G needle only during injection.

### Visual stimulation protocol

To trigger a neuronal response in the mouse brain, we exposed the eyes of head-fixed, lightly anesthetized darkness-habituated mice to a light stimulation protocol (Fig. 2A). The protocol comprised 3 sets of 15 seconds-long blue light flashes (470 nm LED, 3-6 Hz flashing frequency) interleaved with 45 seconds-long periods of darkness (Fig. 2C). This protocol was designed to evoke responses in brain structures that are part of the visual system, namely the visual cortex, the superior colliculus and the lateral geniculate nucleus (LGN) (28). We selected our imaging plane to optimally capture the LGN and assess the subcortical imaging capabilities of enhanced fUS. For each experiment, we first conducted a baseline acquisition without any bolus administration, followed by a second acquisition during which we injected a bolus of either saline, GVs or MBs after a 45-second baseline period. We acquired 4 to 5 datasets for each group (saline, MBs and GVs), each in a separate mouse experiment.

### In vivo data processing

After processing power Doppler images (Fig. 2B), we analyzed the signals recorded 90 seconds after the start of the acquisition, after the initial bolus peak has passed, leaving a relatively stable contrast (Fig. 2D). The Doppler data was denoised in the time domain using a 4-point moving average. To remove the slope of the post-bolus washout, we fitted the Doppler intensity time traces in every pixel with a linear regression model and subtracted the linear trend from the experimental data (Fig. 2E).

Neural activity maps (Fig. 2F) were generated by cross-correlation of the temporal signal in each pixel of the de-trended power Doppler datasets with the hemodynamic response function (HRF) to the stimulus. The HRF was computed based on a typical averaged response over all the voxels showing a response to light with a signal > 3 spatial standard deviations (STDs).

In order to characterize the fast-time temporal fluctuation of the contrast-enhanced Doppler signal during the bolus washout phase (Fig. 3), we fitted the Doppler signal from t=60 s with a double exponential function, which we subtracted from the data in every pixel. Subsequently, we derived the standard deviation of signal using a Gaussian distribution fit, and computed the variance as the square of the standard deviation.

**Figure 3.**
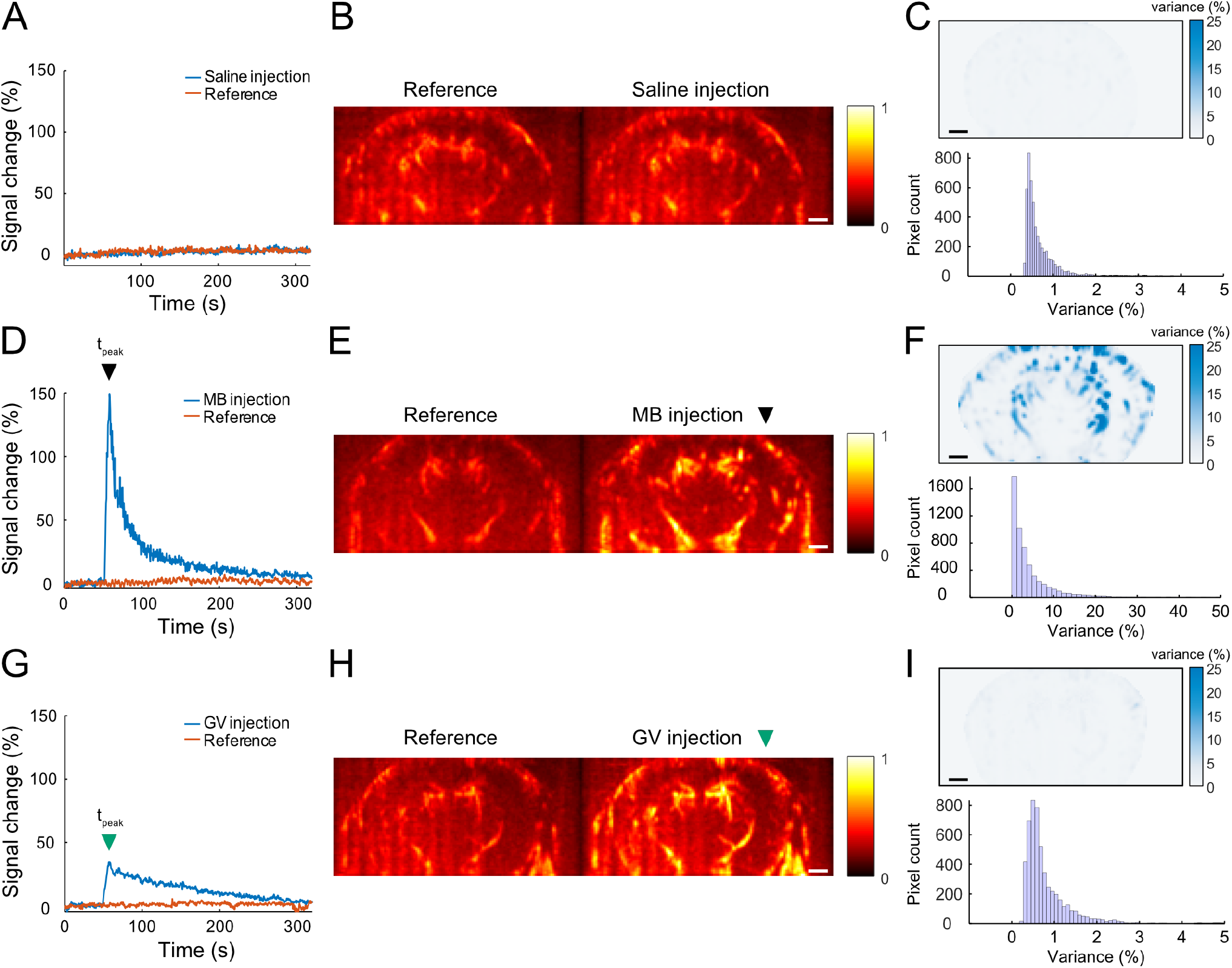
GV enhancement of ultrafast ultrasound Doppler signals. A/ Mean brain signal change over time with and without saline. B/ Power Doppler images at 60s with and without saline. C/ (Top) Variance of the fUS signal fluctuation per pixel in the presence of saline. (Bottom) Histogram of the variance observed in image pixels. D/ Mean brain signal change over time with and without MBs. E/ Power Doppler images at peak enhancement with and without MBs. F/ (Top) Variance of the fUS signal fluctuation per pixel in the presence of MBs. (Bottom) Histogram of the variance observed in image pixels. G/ Mean brain signal change over time with and without GVs. H/ Power Doppler images at peak enhancement with and without GVs. I/ (Top) Variance of the fUS signal fluctuation per pixel in the presence of GVs. (Bottom) Histogram of the variance observed in image pixels. Scale bars represent 1 mm.

We overlaid neural activation maps on top of power Doppler images of the cerebral vasculature (Fig. 4B, D, F) using a threshold of 3 STDs above the noise level (8).

**Figure 4.**
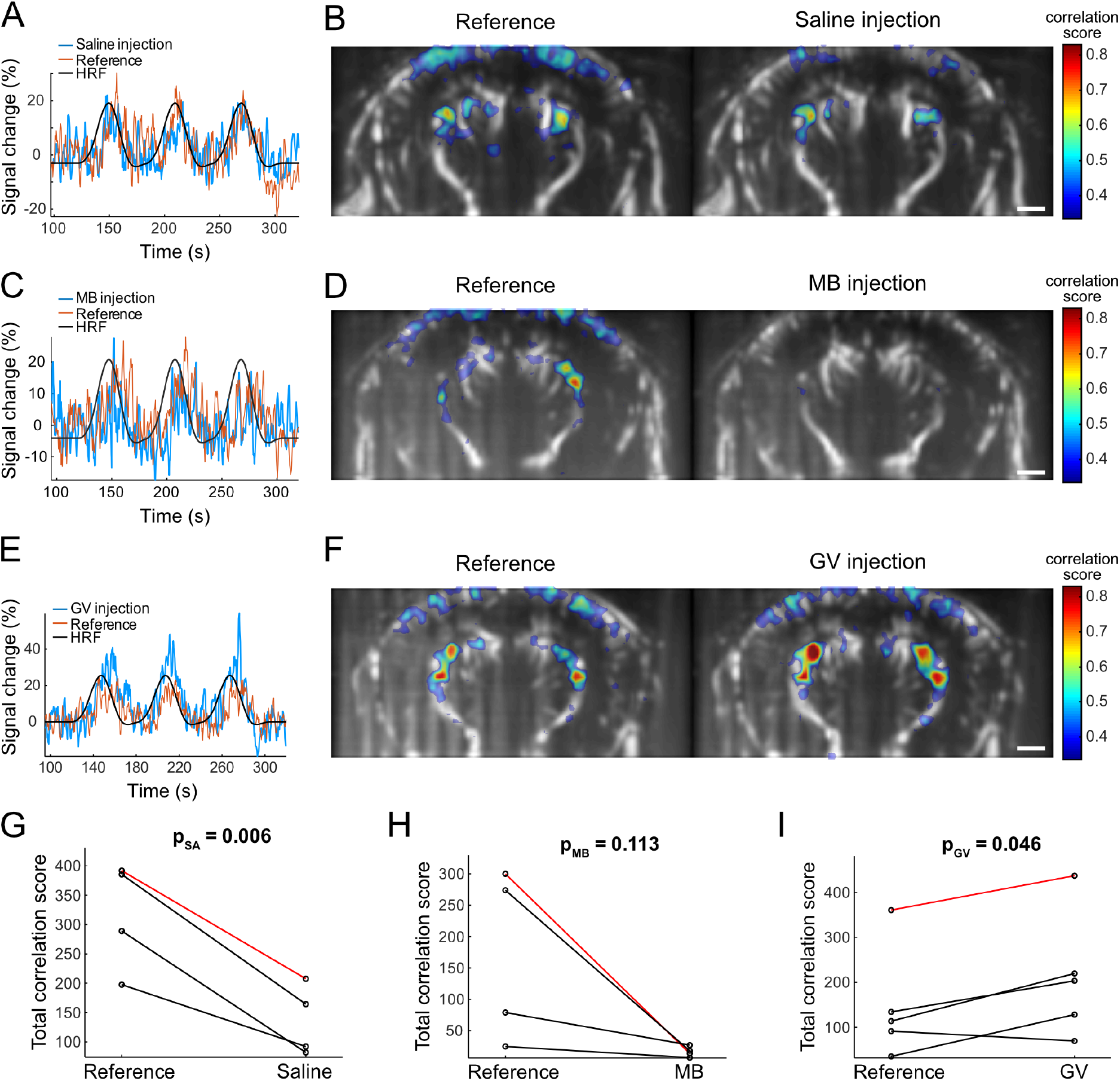
GV enhancement of transcranial fUS signals. A/ C/ E/ fUS signals in the most responsive LGN pixel with and without bolus injection of saline, MBs, and GVs, respectively. B/ D/ F/ Activation maps overlaid on power Doppler images of the mouse brain with and without bolus injection of saline, MBs, and GVs, respectively. G/ H/ I/ Integrated correlation scores (number of pixels times their correlation score) with and without bolus injection of saline, MBs, and GVs, respectively. The points connected by the red lines correspond to the examples reported in B, D and F. Scale bars represent 1 mm.

Finally, we computed integrated correlation scores (Fig. 4G-I) by summing the correlation score of each pixel above 3 STDs within the masked brain image. Statistical significance before and after bolus injection was characterized using a two-tailed paired t-test. All ultrasound images displayed in the manuscript were linearly interpolated by halving intervals in the x and y directions.

### Cerebral blood flow velocity computation

We segmented SVD-filtered beamformed IQ data into discrete frequency bands of 20 Hz, which corresponds to velocity bands of 1 mm/s, using 6th-order Butterworth bandpass filters (10-30 Hz to 50-70 Hz with increments of 10 Hz; and 80-100 Hz to 460-480 Hz with increments of 20 Hz). This processing approach enabled the generation of a set of power Doppler images that map cerebral blood flow in discrete velocity ranges of 1 mm/s (centered on 1 - 4.5 mm/s with increments of 0.5 mm/s, and 5.5 – 23.5 mm/s with increments of 1 mm/s) (29). We subsequently analyzed the distribution of cerebral blood flow velocities in the activated LGN pixels (Fig. 5), that were segmented from the activation maps using a correlation coefficient threshold of 0.6. Finally, we computed the Doppler intensity in each band between during rest and during light-evoked stimulation of the LGN in the absence and presence of GVs.

**Figure 5.**
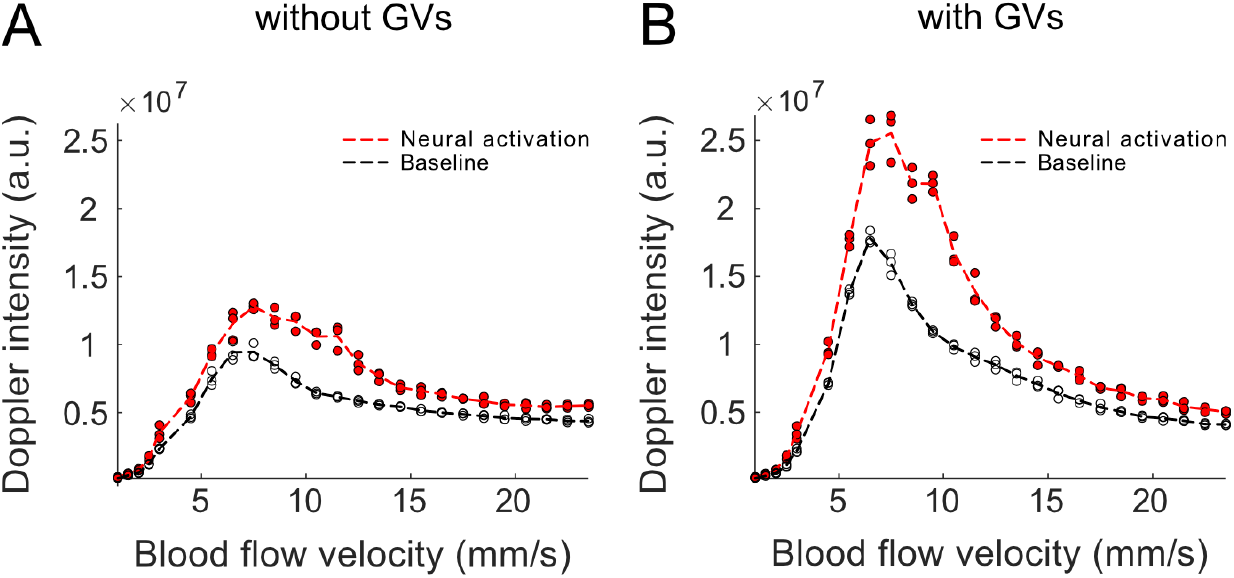
Distribution of cerebral blood flow velocities in the LGN. A/ Linear Doppler intensities in bandpass filtered velocity bins of 1mm/s at baseline (black) and during activation (red) in the absence of GVs. B/ Linear Doppler intensities in bandpass filtered velocity bins of 1mm/s at baseline (black) and during activation (red) in the presence of GVs.

## RESULTS

### GVs enhance Doppler contrast across velocities

*In vitro* results showed that, at a 2 mm/s flow velocity, GVs can sustain ultrasound peak positive pressures ranging from 135 kPa to 507 kPa without showing any decrease in CFR (Fig. 1D). Above 507 kPa, the CFR enhancement of Doppler signals attributed to GVs decreased continuously down to 846 kPa, indicating that most GVs were collapsed at this pressure. In contrast, MBs began to collapse above 254 kPa (Fig. 1D). Being able to withstand higher insonation pressures is beneficial because the intensity of signals backscattered from both the endogenous red blood cells and the contrast agents typically scales as pressure squared.

Our *in vitro* experiments also revealed that GVs are more accurate reporters of flow velocity below 500 μm/s than MBs and DF alone. At 254 kPa (Fig.1E), fluid containing GVs measured the syringe pump flow velocity accurately down to 50 μm/s, while DF itself reflected flow velocity accurately down to 200 μm/s, and MBs where only accurate above 500 μm/s. This may be due to the dominance of Stokes drag over other forces such as buoyancy in the case of GV nanoparticles, while playing a less dominant role for the larger MBs and DF particles. At 592 kPa (Fig. 1F), a partially destructive pressure level for both MBs and GVs, DF particles and MBs were accurate reporters of flow velocities down to 100 μm/s, whereas GVs where accurate reporters down to 50 μm/s flow velocities. The higher MB accuracy at high pressure is likely due to the higher SNR and to the destruction of static bubbles in the channel.

These results establish the basic ability of GVs to enhance Doppler contrast across a wider range of velocities and withstand higher insonation pressures than MBs.

### GVs yield smooth Doppler enhancement in vivo

To compare the *in vivo* performance of MBs and GVs in enhancing power Doppler and fUS signals, we used a setup combining contrast agent injection, transcranial fUS imaging and stimulation of the visual system in anesthetized mice (Fig. 2). We started by evaluating the power Doppler signal enhancement conferred by a single bolus injection of saline, MBs or GVs in the absence of light stimulation (Fig. 3). Key performance parameters included the magnitude of signal enhancement and the temporal noise, which are expected to have opposite impacts on the ability of fUS to detect neural activity. As expected, control injections of saline did not modify the Doppler signal intensity over time compared to the reference acquisition (Fig. 3A). Consequently, transcranial power Doppler images at t = 60 s without and with saline (Fig. 3B) looked similar. For a saline injection, the mean variance of the fast-time fluctuation of the saline-enhanced Doppler signal was 0.66% (Fig. 3C).

The bolus injection of MBs (Fig. 3D) enhanced the Doppler signal significantly, peaking at 149% compared to the pre-injection baseline level at t = 58.5 s. Transcranial power Doppler images at t = 58.5 s without and with MBs (Fig. 3E) revealed a clear enhancement across the entire brain due to MBs. The variance of the signal was characterized in the washout phase using a double exponential fit (R^2^ = 0.97). Following MB injection, the mean variance was 4.45% (Fig. 3F), one order of magnitude above the saline value.

The bolus injection of GVs (Fig. 3G) enhanced the Doppler signal by 34% compared to the pre-injection baseline level at t = 57.5 s. Transcranial power Doppler images at t = 57.5 s without and with GVs (Fig. 3H) revealed a clear enhancement throughout the brain. The variance of the signal was characterized in the washout phase using a double exponential fit (R^2^ = 0.97). For the GV injection, the mean variance was 0.79% (Fig. 3I) – on the same order of magnitude as the saline value, and 5.6 times smaller than for MBs. These results demonstrate the ability of GVs to provide a similar magnitude of pseudo-steady hemodynamic contrast enhancement as MBs, while producing substantially less signal fluctuation.

### GVs enhance transcranial fUS signals in mice

Next, we evaluated the enhancement of visually evoked fUS signals in the LGN of anesthetized mice after a single bolus injection of saline, MBs or GVs. fUS contrast was clearly observed for both the reference measurement (blood contrast alone) and after saline injection (Fig. 4A, B). In the single-trial recording shown in Fig. 4A, the peak fUS signal activation in the masked brain was 30% and 22%, without and with saline, respectively. In functional activation maps (Fig. 4B), the highest LGN correlation scores were 0.66 and 0.65, without and with saline injection, respectively. We observed that saline did slightly decrease correlation scores in activation maps, especially in the visual cortex. This may be due to the local dilution of the red blood cell concentration in the dense capillary networks of the cortex.

In the single-trial recording reported in Fig. 4C, the peak fUS signal activation of the reference acquisition was 27%, and 28% with MBs injected. However, the MB-enhanced fUS signal showed substantially larger fluctuations than the reference signal. As a result, in correlation maps (Fig. 4D), the highest LGN correlation score dropped from a pre-injection value of 0.70 to only 0.37 after MB injection, indicating that the statistical power to detect fUS signals resulting from neural activity diminished due to the large MB-induced signal fluctuation.

In contrast, GVs produced enhanced fUS responses and statistical correlation. In the single-trial recording reported in Fig. 4E, the peak fUS signal activation of the reference acquisition was 25%, going up to 60% after GV injection. In correlation maps (Fig. 4F), the highest LGN correlation score in the reference acquisition was 0.75, and increased to 0.84 with GVs administered, indicating a clear enhancement of functional signals.

Overall findings in groups of N≥4 mice indicated a statistically significant increase in the integrated correlation score of fUS-recorded LGN activations with GVs administered compared to red blood cells alone (p-value = 0.046) (Fig. 4I). In contrast, saline and MB injections appeared to degrade fUS-recorded LGN activations compared to red blood cells contrast (p-value = 0.006 and 0.113) (Fig. 4G, H).

To further characterize the impact of GV administration on the information content of hemodynamic fUS signals, we analyzed contrast enhancement as a function of the observed blood flow velocities in activated patches of the LGN by segmenting Doppler images into velocity bands of 1 mm/s (29). Fig. 5 reports the blood flow velocity profiles in the LGN (in pixels with correlation scores above 0.6) for the data set shown in Fig. 4F. In the absence of GVs (Fig. 5A), the cerebral blood flow velocity profile during light-evoked LGN activation revealed an overall increase of the Doppler intensity across all the cerebral blood flow velocities sampled (1 mm/s to 24 mm/s) compared to the cerebral blood flow velocity profile at rest. In addition, a noticeable increase in Doppler intensity was observed around 10 mm/s. In the presence of GVs (Fig. 5B), Doppler intensities were increased, and the enhancement did not significantly bias the cerebral blood flow velocity profiles, which remained similar to Fig. 5A. This indicates that GVs enhance all classes of vessels contributing to the fUS signals.

## DISCUSSION

Together, our results establish the potential of GVs to serve as hemodynamic enhancers for functional ultrasound imaging. Our *in vitro* flow phantom experiments confirmed the hypothesis that GVs can enhance ultrasound Doppler signals on the same order of magnitude in a pseudo-steady state as MBs, but sustain higher ultrasound pressures (507 kPa versus 254 kPa) and more accurately report low flow velocities (down to 50 μm/s). The first difference is of importance for ultrasound imaging because higher pressure transmission leads to a higher dynamic range in Doppler images. The second finding is critical for future efforts to extend fUS imaging to include the capillary level of cerebral vasculature, a vascular compartment which is currently poorly sensed by this technique due to limited sensitivity below a few mm/s (2), whereas capillary flows extend below 1 mm/s (30).

Our *in vivo* results demonstrate that GVs provide similar pseudo-steady enhancement of Doppler signals compared to MBs (despite a lower peak enhancement), without introducing additional temporal fluctuation into the Doppler signal. This smooth enhancement leads to a more effective functional signal amplification using GVs than MBs. This finding is consistent with the fact that GV nanostructures are relatively numerous, monodisperse and well-embedded in blood flow. In contrast, commercial MBs are polydisperse micron-scale contrast agents that respond acoustically in a non-coherent way.

To enable longitudinal studies using GVs as intravascular fUS enhancers, the circulation time of GVs will need to be extended, for example using surface modifications such as PEGylation (31). In addition, the ability to engineer GV properties at the genetic level may enable the optimization of GVs for Doppler contrast. Finally, future studies could investigate the use of engineered GVs that exhibit nonlinear scattering (21, 24, 32) to increase the specificity and resolution of GV-enhanced functional ultrasound imaging.

## CONCLUSION

Taken together, our results demonstrate that GVs provide superior performance as enhancers of fUS compared to MBs due to GVs’ ability to withstand higher pressure, more faithfully report slower flow velocities, and not increase the temporal fluctuation of the measurement. Further engineering for enhanced circulation time and brighter contrast would make GVs a preferred enhancer for fUS imaging.

## ACKNOWLEDGEMENTS

We thank Di Wu and Thomas Deffieux for helpful discussions. DM is supported by a Human Frontiers Science Program Cross-Disciplinary Postdoctoral Fellowship (Award No. LT000637/2016). This research was funded by the National Institutes of Health (grant U01NS099724 to MGS). Related research in the Shapiro laboratory is also supported by the Heritage Medical Research Institute, Burroughs Wellcome Career Award at the Scientific Interface, the Pew Scholarship in the Biomedical Sciences and the Packard Fellowship for Science and Engineering.

